# Site level factors that affect the rate of adaptive evolution in humans and chimpanzees; the effect of contracting population size

**DOI:** 10.1101/2021.05.28.446098

**Authors:** Vivak Soni, Ana Filipa Moutinho, Adam Eyre-Walker

**Affiliations:** School of Life Sciences, University of Sussex, Brighton, BN1 9QG; Department for Evolutionary Genetics, Max Planck Institute for Evolutionary Biology, Plon, Germany

## Abstract

It has previously been shown in other species that the rate of adaptive evolution is higher at sites that are more exposed in a protein structure and lower between amino acid pairs that are more dissimilar. We have investigated whether these patterns are found in the divergence between humans and chimpanzees using an extension of the MacDonald-Kreitman test. We confirm previous findings and find that the rate of adaptive evolution, relative to the rate of mutation, is higher for more exposed amino acids, lower for amino acid pairs that are more dissimilar in terms of their polarity, volume and lower for amino acid pairs that are subject to stronger purifying selection, as measured by the ratio of the numbers of non-synonymous to synonymous polymorphisms (p_N_ /p_S_). However, the slope of this latter relationship is significantly shallower than in Drosophila species. We suggest that this is due to the population contraction that has occurred since humans and chimpanzees diverged. We demonstrate theoretically that population size reduction can generate an artefactual positive correlation between the rate of adaptive evolution and any factor that is correlated to the mean strength of selection acting against deleterious mutations, even if there has been no adaptive evolution (the converse is also expected). Our measure of selective constraint, p_N_ /p_S_, is negatively correlated to the mean strength of selection, and hence we would expect the correlation between the rate of adaptive evolution to also be negatively correlated to p_N_ /p_S_, if there is no adaptive evolution. The fact that our rate of adaptive evolution is positively correlated to p_N_ /p_S_ suggests that the correlation does genuinely exist, but that is has been attenuated by population size contraction.

## Introduction

The rate of adaptive evolution in protein coding genes varies at several different levels. First, the rate of adaptive evolution appears to differ between species. Some species, including many plants (Bustamante et al. 2002; Barrier et al. 2003; Schmid et al. 2005; Gossman et al. 2010; also see Strasburg et al. 2009; Ingvarsson et al. 2010; Slotte et al. 2010) and the yeasts of the genus Saccharomyces (Gossman et al. 2012), appear to go through very little adaptive evolution, whilst many other species, including Drosophilids (Smith and Eyre-Walker 2002; Sawyer et al. 2003; Eyre-Walker and Keightley 2009; Haddrill et al. 2010), rodents (Halligan et al. 2010) and many multicellular animals (Galtier 2016; Rousselle et al. 2019), go through extensive adaptive evolution. The reasons for this variation remain unclear. It has been suggested that population size might be a factor; if adaptation is mutation limited, then one might expect species with large population sizes to adapt faster because they will generate the required mutation faster. There is some evidence that species with large population sizes undergo significantly faster adaptive evolution (Gossman et al. 2012; Bataillon et al. 2015; Corbett-Detig et al. 2015; Rousselle et al. 2019), though in Galtier (2016) the correlation with ω_a_ is non-significant. Furthermore, it is unclear whether species are ever limited by the supply of mutations -there appears to be abundant genetic variation for most traits -and even if they are limited, species with large population sizes are predicted to be closer to their optimal fitness, and hence they may not have to adapt as much as species with small population sizes (Lourenco et al. 2013).

At the next level down, there appears variation in the rate of adaptation between genes. This is in part due to differences in function, with genes involved in immunity (Clark et al. 2003; Nielsen et al. 2005; Chimpanzee Sequencing and Analysis Consortium, 2005; Sackton et al. 2007; Obbard et al. 2009), interaction with viruses (Enard, et al. 2016) and male reproductive success (Proschel et al. 2006; Haerty et al. 2007) having high rates of adaptive evolution. Other factors also seem to be important, with the rate of adaptive evolution being higher in genes that recombine frequently (Presgraves, 2005; Betancourt et al. 2009; Arguello et al. 2010; Mackay et al 2012; Campos et al. 2014; Castellano et al. 2016; Moutinho et al. 2019), are located in regions of the genome with low functional DNA density (Castellano et al. 2016), have high mutation rates (Castellano et al. 2016) and reside on the X-chromosome (MacKay et al. 2012; Langley et al. 2012; Campos et al. 2014). Genes that have lower expression levels (Pal et al. 2001; Subramanian and Kumar, 2004; Wright et al. 2004; Rocha and Danchin, 2004; Lemos et al. 2005) or shorter coding sequence length (Zhang, 2000; Lipman et al. 2002; Liao et al. 2006), also seem to have higher rates of adaptation.

Finally, there appears to be variation at the site level. This variation has been widely documented in site-level tests that compare the rate of non-synonymous to synonymous substitution (for example, Liberles et al. 2012). A number of factors seem to affect rates of adaptive evolution at the site level including protein secondary structure (Goldman et al. 1998; Guo et al. 2004; Choi et al. 2006) and the relative solvent accessibility (RSA) (Goldman et al. 1998; Choi et al. 2007; Lin et al. 2007; Franzosa and Xia 2009); RSA is a measure of how buried an amino acid is. In both *Drosophila* and *Arabidopsis* species, the rate of adaptive non-synonymous substitution is positively correlated to the relative solvent accessibility (RSA) (Moutinho et al. 2019). This suggests that amino acids on the surface of a protein have higher rates of adaptive substitution than those that are buried (Perutz et al. 1965; Overington et al. 1992; Goldman et al. 1998; Bustamante et al. 2000; Dean et al. 2002; Choi et al. 2006; Lin et al. 2007; Conant and Stadler 2009; Franzosa and Xia 2009; Ramsey et al. 2011). It has also been shown that amino acids that differ strongly in their physio-chemical properties, have lower rates of adaptive evolution than those that are more similar (Bergman and Eyre-Walker, 2019; though see Gojobori et al. 2007 and Chen et al. 2019). Finally, Bergman and Eyre-Walker (2019) also showed that amino acids pairs that are subject to high levels of negative selection have lower rates of adaptive substitution; they measured the level of negative selection using the ratio of the number of non-synonymous to synonymous polymorphisms, p_N_ /p_S_ .

In our analysis we consider whether the rate of adaptive evolution between humans and chimpanzees is correlated to several site level factors previously shown to be particularly important in other species -RSA and various measures of the difference between amino acids, and the overall level of negative selection acting on amino acid pairs. We find negative correlations between the rate of adaptive evolution and the difference in amino acid physio-chemical properties, and a positive correlation between the rate of adaptive evolution and RSA and our measure of negative selection.

## Results

We set out to investigate whether several site-level factors affect the rate of adaptive and non-adaptive evolution in hominids – relative solvent accessibility (RSA), and measures of physio-chemical (volume and polarity) and the level of negative selection acting on mutations between two amino acids (p_N_ /p_S_). We measure the rates of adaptive and non-adaptive evolution using the statistics ω_a_ and ω_na_, which are respectively estimates of the rate of adaptive and non-adaptive evolution relative to the mutation rate. Both statistics were estimated using an extension of the McDonald-Kreitman method (McDonald and Kreitman, 1991), in which the pattern of substitution and polymorphism at neutral and selected sites is used to infer the rates of substitution, taking into account the influence of slightly deleterious mutations. We use the method implemented in GRAPES (Galtier, 2016), which is a maximum likelihood implementation of the second method proposed by Eyre-Walker and Keightley (2009).

### Relative solvent accessibility

Previous studies have shown that amino acid residues at the surface of proteins evolve faster than those at the core (Goldman et al. 1998; Choi et al. 2006; Lin et al. 2007; Franzosa and Xia, 2009). These studies do not distinguish whether this higher substitution rate is due to reduced selective constraints on exposed residues or an increased rate of adaptive substitutions (or both). Moutinho et al (2019) disentangled these effects by estimating both the rates of adaptive and non-adaptive evolution across several RSA categories in *Drosophila* and *Arabidopsis*, finding positive correlations between RSA and the rates of both adaptive and non-adaptive substitution. Their findings suggest that both reduced negative selection and a higher rate of adaptive evolution operate on more exposed residues. We find a significant correlation between the rate of adaptive evolution and RSA (r=0.486, p<0.001) when we use a weighting by the reciprocal of the variance of the rate of adaptive or non-adaptive evolution. However, the correlation with the rate of non-adaptive evolution is non-significant (r=0.001, p=0.324) in hominids (figure 1).

**Figure 1:**
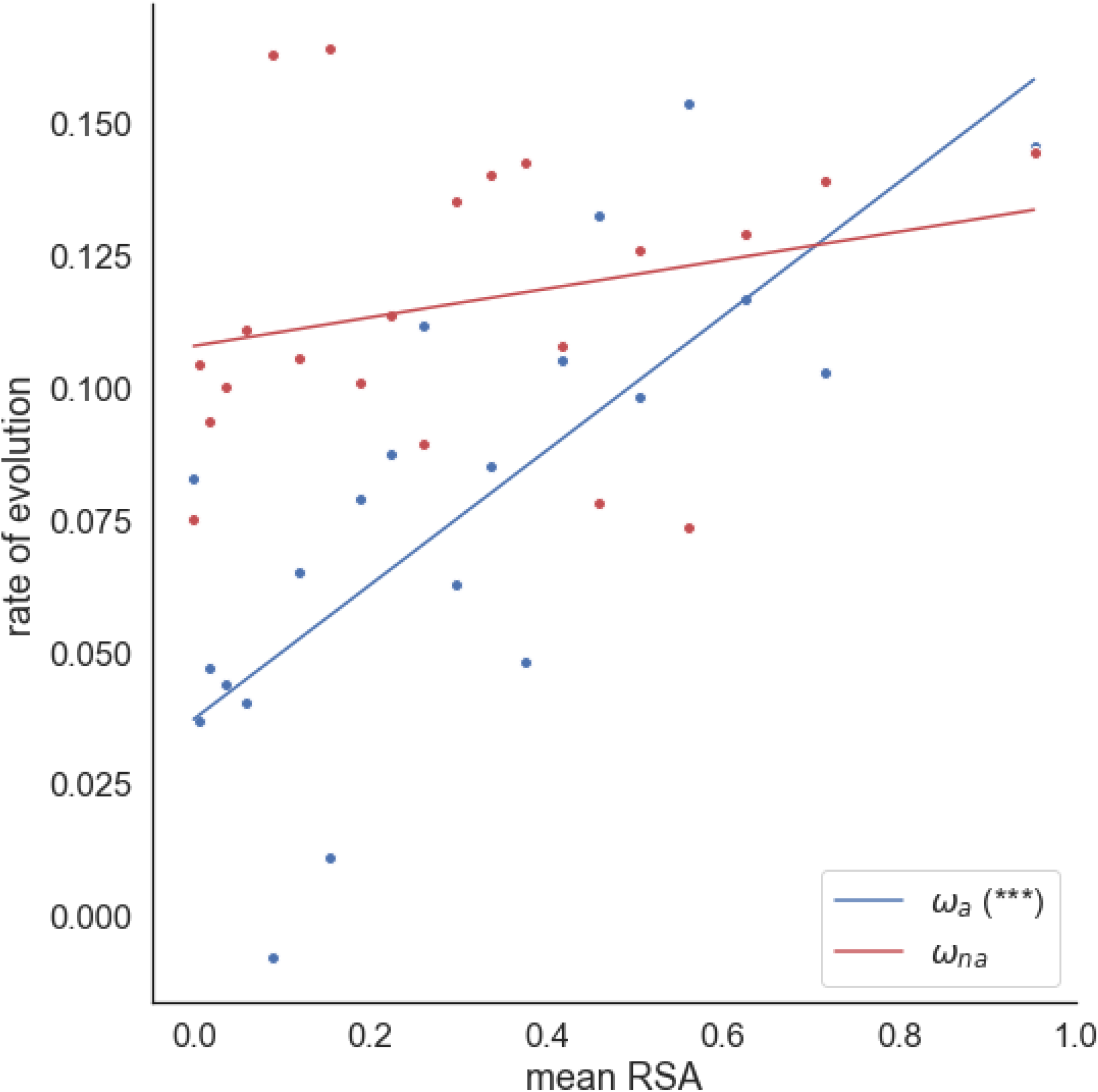
Estimates of ω_a_ and ω_na_ plotted against mean relative solvent accessibility. Data binned into 20 RSA bins of roughly equal number of sites. For each analysis, a weighted linear regression is fitted to the data. The respective significance of each correlation is shown in the plot legend (*P < 0.05; **P < 0.01; ***P < 0.001; “.” 0.05 ≤ P < 0.10) for ω_a_ and ω_na_). Regression is weighted by the reciprocal of the variance for each estimate of ω_a_ and ω_na_, which were estimated by bootstrapping the data by gene 100 times for each data point.

### Amino acid dissimilarity

To investigate whether the rates of adaptive and non-adaptive evolution are affected by amino acid dissimilarity, we estimated ω_a_ and ω_na_ between all 75 pairs of amino acids that are separated by a single mutational step in hominids. Bergman and Eyre-Walker (2019) found negative correlations between measures of amino acid dissimilarity (differences in volume and polarity) and ω_a_ between *Drosophila* species. We find that the rate of adaptive substitution is significantly negatively correlated to the difference in volume (r = −0.290, p = 0.018) and polarity (r = −0.269, p = 0.027) (figures 2a and 2b) when we fit a weighted linear regression to the data, suggesting that the rate of adaptive evolution is higher between more physiochemically similar amino acids. Similar negative correlations are observed for the rate of non-adaptive evolution (volume difference: r = −0.545, p < 0.001; polarity difference: r = −0.170, p<0.001). The slopes are significantly steeper for ω_na_ (Table 1); however, this appears to be simply because rates of non-adaptive evolution are greater than rates of adaptive evolution; when we divide ω_a_ and ω_na_ by their means, the slopes are not significantly different.

**Figure 2:**
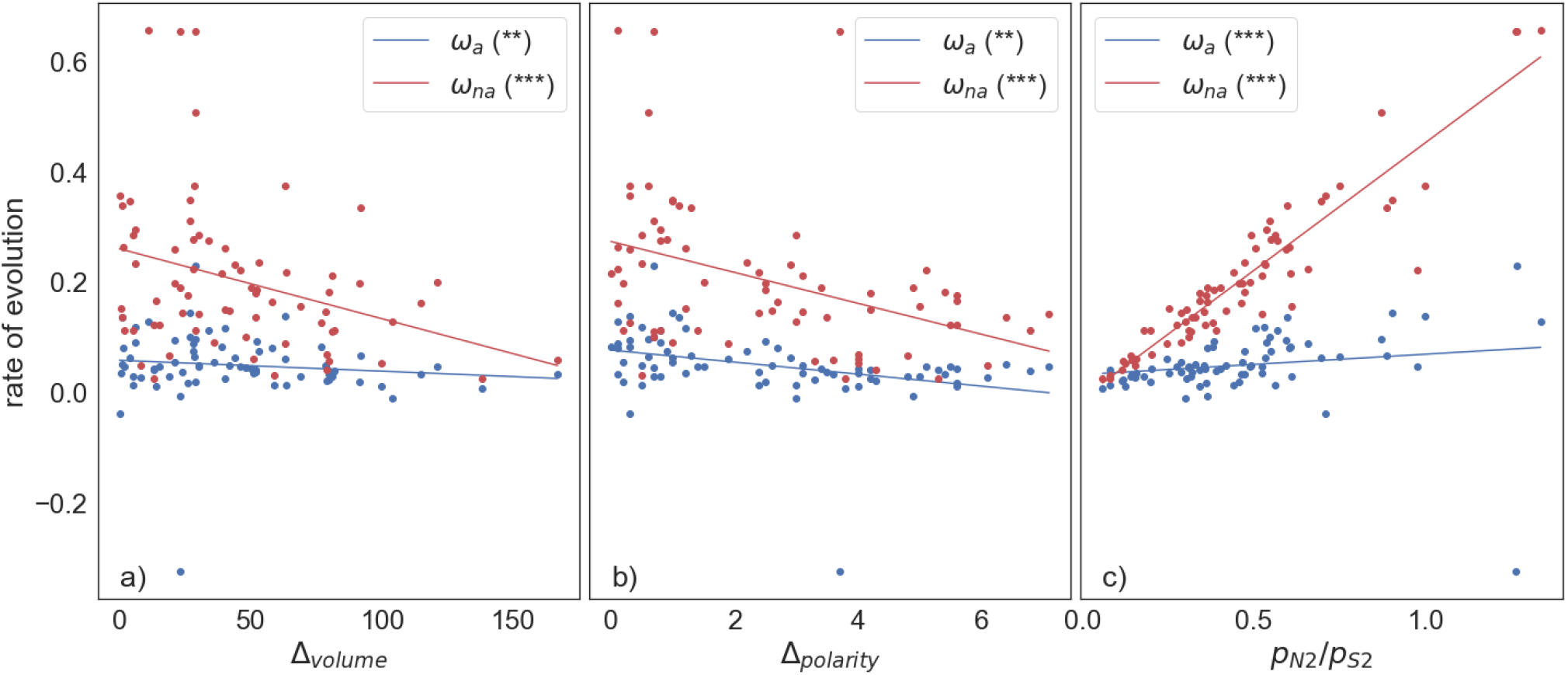
The adaptive and non-adaptive substitution rate plotted against the difference in a) volume, b) polarity and c) the ratio of nonsynonymous to synonymous polymorphisms, p_N2_ /p_S2_ . In c) the polymorphisms are split by sampling from a hypergeometric distribution, with one set used to calculate rates of adaptive and non-adaptive substitution and the other to estimate the two polymorphism statistics. A weighted linear regression is fitted to the data, weighted by the variance of each estimate. The respective significance of each correlation is shown in the legend (*P < 0.05; **P < 0.01; ***P < 0.001; “.” 0.05 ≤ P < 0.10).

**Table 1.** The slope of the relationship between ω_a_ and ω_na_ and the Δvolume and Δpolarity; rescaled values are where ω_a_ and ω_na_ have been divided by their means. Significance was measured using analysis of variance.

| Statistic | Rescaled | $\omega_a$ | | $\omega_{na}$ | | Sig. |
| --- | --- | --- | --- | --- | --- | --- |
|  |  | Slope | SE | Slope | SE |  |
| Δvolume | No | -0.00027 | 0.000098 | -0.0010 | 0.00026 | 0.012 |
| Δpolarity | No | -0.0064 | 0.0020 | -0.022 | 0.0054 | 0.010 |
| Δvolume | Yes | -0.0054 | 0.0020 | -0.0051 | 0.0013 | n.s. |
| Δpolarity | Yes | -0.13 | 0.042 | -0.11 | 0.027 | n.s. |

The difference in polarity and volume are not significantly correlated to each other (r=0.122, p=0.258), so it seems likely that both volume and polarity have an influence over the rate of adaptive and non-adaptive evolution. A multiple regression confirms this for ω_na_ with both factors being highly significant and of similar influence, as judged by standardised regression coefficients (Δvolume b_s_ = −0.29, p = 0.015; Δpolarity b_s_ = −0.31, p = 0.008). For ω_a_, only Δpolarity is significant (Δvolume b_s_ = −0.19, p = 0.14; Δpolarity b_s_ = −0.27, p = 0.036); the loss of significance for Δvolume is probably due to a loss of power due to lack of data; in multiple regression we are effectively holding one variable constant and testing whether the other remains significant.

Volume and polarity reflect only two of the multiple ways in which amino acids differ. As an alternative measure of amino acid dissimilarity Bergman and Eyre-Walker (2019) suggest using the ratio of non-synonymous to synonymous polymorphism; p_N_ /p_S_ is expected to decrease as the strength of selection against deleterious mutations increases. We find that hominids are consistent with this expectation as p_N_ /p_S_ is negatively correlated with both amino acid volume difference (r = −0.456, p < 0.001) and polarity difference (r = −0.269, p = 0.047) (supplementary figure S1). Polymorphism data is used to estimate both the rates of adaptive and non-adaptive substitution, meaning that p_N_ /p_S_ is not statistically independent of either measure. To account for this source of sampling error we follow the method of Bergman and Eyre-Walker (2019), resampling the site frequency spectrum using a hypergeometric distribution to generate two independent spectra. One of these is used to estimate p_N_ /p_S_ (referred to as p_N2_ /p_S2_) and the other is used to estimate ω_a_ and ω_na_, therefore removing the nonindependence between p_N_ /p_S_ and ω_a_ and ω_na_ . We find that ω_a_ is positively correlated to p_N2_ /p_S2_ (r = 0.419, p<0.001) in hominids, consistent with previous findings in *Drosophila* (Bergman and Eyre-Walker, 2019). Consistent with our physicochemical dissimilarity correlations, ω_na_ is also shows a positive correlation with p_N_ /p_S_, but a stronger one (r = 0.882, p < 0.001) (figure 2c).

It is possible that the correlations between ω_a_ and ω_na_ and various site level factors are interrelated; for example, the positive correlation between ω_a_ and RSA might be due to amino acids that are found exposed on the surface of proteins being one mutational step closer to similar amino acids. However, there is no correlation between the average RSA of an amino acid and the average difference in volume or polarity to its one mutation step neighbours (RSA-volume: r=-0.171, p=0.471; RSA-polarity: r=0.059, p=0.803 – supplementary figure S1).

### Biased gene conversion

Biased gene conversion can potentially impact estimates of the rate of adaptive evolution, since it increases the fixation probability of Weak (W) to Strong (S) alleles relative to S>W neutral alleles, more than it increases levels of W>S polymorphisms relative to S>W polymorphisms; a problem exacerbated by differences in base composition between synonymous and non-synonymous sites (Galtier and Duret, 2007; Berglund et al. 2009; Ratnakumar et al. 2010; Rousselle et al. 2020). To investigate whether the correlation between the rates of adaptive and non-adaptive evolution and our measures of amino acid dissimilarity are due to BGC we restricted the analysis to polymorphisms and substitutions that involve nucleotide changes that are unaffected by BGC – i.e. A<>T and G<>C changes. This reduces our dataset substantially removing 80% of our substitutions and polymorphisms, and reducing the amino acid analysis to just 12 amino acid pairs. However we find that the correlations between ω_a_, RSA, difference in volume and p_N_ /p_S_ all remain significant (RSA: r = 0.260, p < 0.05; volume difference: r=-0.576, p<0.01; polarity difference: r=-0.166, p<0.1; p_N2_ /p_S2_ : r =0.796, p<0.001); the correlations between ω_na_ and the difference in volume and p_N2_ /p_S2_ remain significant (RSA: r=0.011, p=0.370; volume difference: r=0.513, p<0.01; polarity difference: r=0.115, p=0.150; p_N2_ /p_S2_ : r=0.804, p<0.001).

## Discussion

Our analyses showed that the rate of adaptive evolution is significantly positively correlated to the relative solvent accessibility and measures of amino acid dissimilarity in humans and chimpanzees, being consistent with previous findings in *Drosophila* (Moutinho et al. 2019; Bergman and Eyre-Walker, 2019). We find similar correlations for the rate of non-adaptive evolution, except in the case of RSA. The majority of these correlations are robust to controlling for the effects of biased gene conversion by restricting the analysis to mutational changes that are unaffected by BGC, despite the huge reduction in the dataset.

It is of interest to ask how the slopes of the relationships between ω_a_ and the site level factors compare to those previously estimated in Drosophila species. We find that the slope is not significantly different for RSA, the difference in volume and the difference in polarity. However, there is a very striking difference in the slope between ω_a_ and p_N_ /p_S_ between hominids and Drosophilids (Table 2). This might be because of population size contraction, which is believed to have occurred since the common ancestor of humans and chimpanzees along both lineages (Holboth et al. 2007; Burgess and Yang 2008; Prado-Martinez et al. 2013; Schrago 2014). It is well known that population size expansion leads to an overestimate, and population size contraction to an underestimation of the rate of adaptive evolution in MK-style approaches (McDonald and Kreitman 1991; Eyre-Walker 2002). This is because the effective population differs between the divergence and polymorphism phases of evolution if the population size changes;. If there has been population size contraction, there will be an excess of slightly deleterious mutations segregating in the polymorphism data because they had no chance of being fixed during the divergence phase with its larger effective population size. How do population size expansion and contraction affect the relationship between ω_a_ and other variables?

**Table 2.** Slopes of the regressions between ω_a_ and measures of amino acid dissimilarity in hominid and *Drosophila* datasets. In the rescaled analyses, the ω_a_ values have been divided by their mean. The slopes for the *Drosophila* analysis were obtained from the results supplied by Bergman and Eyre-Walker (2019).

| Dataset | Independent variable | Hominids (this analysis) |  | Drosophila (Bergman and Eyre-Walker 2019) |  | Sig. |
| --- | --- | --- | --- | --- | --- | --- |
|  |  | Slope | SE of slope | Slope | SE of slope |  |
| Original | RSA | 0.13 | 0.029 | 0.078 | 0.0065 | n.s. |
| Original | $\Delta\text{Vol}$ | -0.00026 | 0.00010 | -0.00027 | 0.000061 | n.s. |
| Original | $\Delta\text{Pol}$ | -0.0064 | 0.0020 | -0.0047 | 0.0011 | n.s. |
| Original | $p_N/p_S$ | 0.061 | 0.019 | 0.29 | 0.029 | <0.001 |
| Rescaled | RSA | 1.6 | 0.36 | 1.6 | 0.13 | n.s. |
| Rescaled | $\Delta\text{Vol}$ | -0.0054 | 0.0020 | -0.011 | 0.0024 | n.s. |
| Rescaled | $\Delta\text{Pol}$ | -0.13 | 0.042 | -0.18 | 0.041 | n.s. |
| Rescaled | $p_N/p_S$ | 1.3 | 0.40 | 11 | 1.1 | <0.001 |

Let us assume that synonymous mutations are neutral and non-synonymous mutations are neutral or subject to selection. The ratio of the non-synonymous to synonymous substitution rates, *ω* = *ω*_*a*_ + *ω*_*na*_ where *ω*_*a*_ and *ω*_*na*_ are the rate of adaptive and non-adaptive non-synonymous substitution relative to the rate of synonymous substitution, which is an estimate of the mutation rate. Hence,

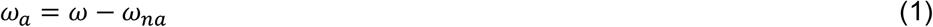

If we assume that all non-synonymous are deleterious with effects drawn from a gamma distribution then

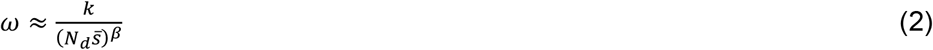

(Welch et al. 2008) where *N*_*d*_ is the effective population size during the divergence phase, *k* is a constant, *β* is the shape parameter of the gamma distribution and 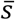 is the mean strength of selection acting against deleterious mutations.

We can also write a simple expression for *ω*_*na*_ . This is estimated in MK type approaches from polymorphism data, using the site frequency spectra (SFS) at synonymous and non-synonymous sites, to estimate the distribution of fitness effects (DFE) at non-synonymous sites. This DFE is then used to infer *ω*_*na*_ . Hence

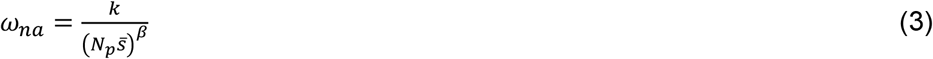

where *N*_*P*_ is the effective population size pertaining to the polymorphism data.

Substituting equation 2 and 3 into 1 we have

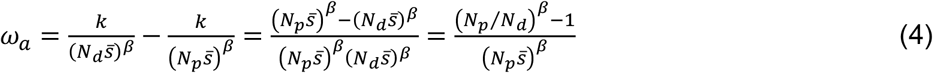

This equation demonstrates that ω_a_ >0 if *N*_*p*_ *>N*_*d*_, and ω_a_ <0 if *N*_*p*_ *<N*_*d*_ as we expect. However, of more interest is the fact that the over-or under-estimation of ω_a_ depends on 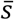, the mean strength of selection acting against deleterious mutations. With population size expansion we predict that ω_a_ will be overestimated but that the magnitude of this overestimation will decrease as the mean strength of selection increases. Conversely, with population size contraction ω_a_ will be under-estimated and this underestimation will diminish as the mean strength of selection increases. Hence, under population size expansion we expect a negative correlation between *ω*_*a*_ and any variable that is correlated to the mean strength of selection and a positive correlation with population contraction, if there is no adaptive evolution.

If we note that

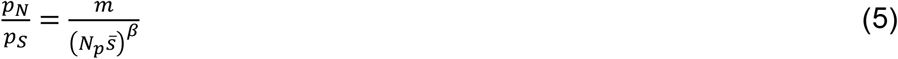

(Welch et a. 2006), where *m* is a constant which depends on how many chromosomes have been sampled, then equation 4 can be rewritten as

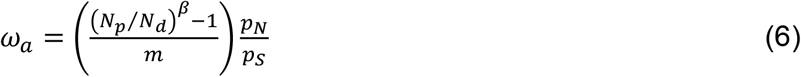

Hence, we expect *ω*_*a*_ to be positively and linearly correlated to p_N_ /p_S_ if there was been population size expansion and negatively correlated if there has been contraction, if there is no adaptive evolution occurring.

The method we have used to estimate *ω*_*a*_ generates an estimate of the mean strength of selection acting against deleterious mutations. We find that 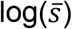 is significantly positively correlated to the difference in volume (r=0.205, p=0.08) and the difference in polarity (r=0.310, p=0.008) and significantly negatively correlated to p_N_ /p_S_ (r=-0.880, p<0.001) but there is no correlation with RSA (r=-0.088, p=0.704) (Figure 3). However, the slope of the relationship is much steeper for p_N_ /p_S_ : if we standardise the variables by subtracting the mean and dividing by the standard deviation the slopes are: RSA = −0.101, Volume b = 0.862, Polarity, b= 1.30, p_N_ /p_S_ = −3.90. Hence, we expect the effects of population contraction to have more of an effect on the relationship between ω_*a*_ and p_N_ /p_S_ than any other variable. This is what we observe: the relationship between *ω*_*a*_ and RSA, volume and polarity is similar in hominids and *Drosophila*, but the relationship between *ω*_*a*_ and pN/pS is much weaker in hominids than *Drosophila* (Table 2).

**Figure 3:**
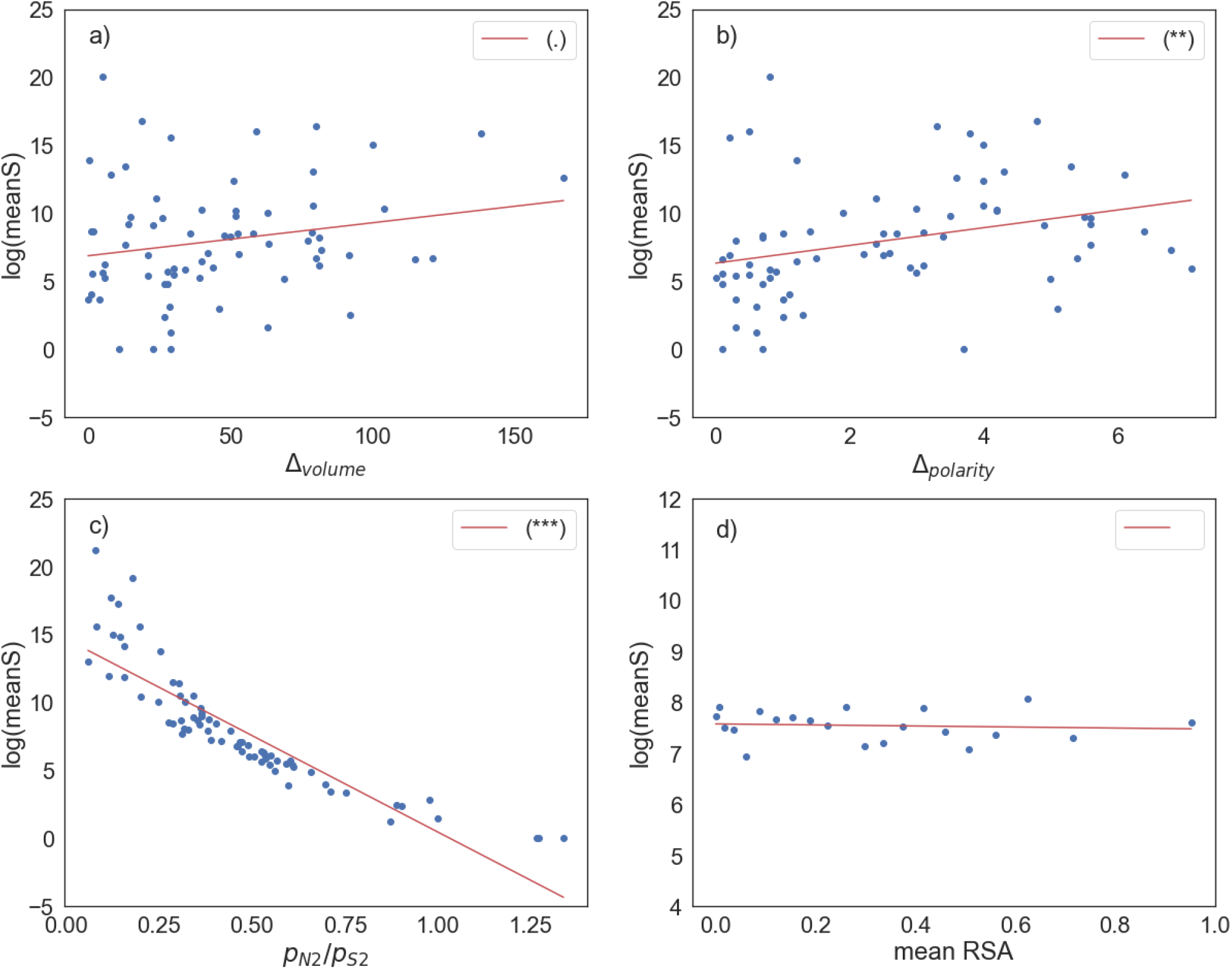
log(meanS) plotted against a) volume difference, b) polarity difference, c) p_N2_ /p_S2_, d) mean RSA. The respective significance of each correlation is shown in the plot legend, (*P < 0.05; **P < 0.01; ***P < 0.001; “.” 0.05 ≤ P < 0.10) based on an unweighted regression fit to the data.

The fact that *ω*_*a*_ is negatively correlated to the difference in volume and polarity, and positively correlated to RSA and p_N_ /p_S_ in a pair of species for which we have independent evidence of population contraction (Holboth et al. 2007; Burgess and Yang 2008; Prado-Martinez et al. 2013; Schrago, 2014) suggests that these correlations are due to variation in the rate of adaptive evolution, because population contraction should lead to the opposite patterns if there is no adaptive evolution.

It is well known that population contraction leads to an underestimate of the rate of adaptive evolution when using MK-style methods (McDonald and Kreitman 1990; Eyre-Walker 2002). Zhen et al. (2021) have argued that the rate of adaptive evolution between humans and chimpanzees has been underestimated, and that they have undergone higher rates of adaptive evolution than Drosophila species. In fact, the average of *ω*_*a*_ across amino acid pairs is significantly higher in hominids than Drosophila (hominids, mean *ω*_*a*_ = 0.0488 (SE = 0.0072; Drosophila mean *ω*_*a*_ = 0.0258 (SE = 0.0024); t-test t = 3.01, p<0.001), so hominids seem to be adapting faster relative to the mutation rate even without taking into account population contraction. What is perhaps surprising is that *ω*_*a*_ is not negative even when we correlate it against factors that appear to influence it. The observed value of *ω*_*a*_ is expectedto be equal to

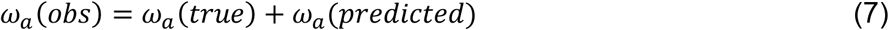

Where *ω*_*a*_ (*true*) is the true value, and *ω*_*a*_ (*predicted*) is the value predicted in the absence of adaptive evolution from equation 4 or 6; i.e. it is the bias in the estimate due to the differences in the effective population size between the divergence and polymorphism phases. For example, *ω*_*a*_ is positively correlated to RSA, however, even those sites with very low RSA, have a positive estimate of *ωa*.This seems surprising and suggests that adaptive evolution is more prevalent than we thought in hominids. However, predicting how much is difficult because we do not know how the effective population size has changed during the divergence of humans and chimpanzees.

We confirm the findings of Moutinho et al. (2019) with respect to RSA -more exposed amino acid residues have higher rates of adaptive evolution. Moutinho et al. (2019) also showed that the rate of non-adaptive evolution is positively correlated to RSA. These observations are consistent with two models of evolution; either the fitness landscape is relatively flat for more exposed residues, or the mutational steps are relatively small. It is difficult to differentiate between these models.

We also confirm the results of Bergman and Eyre-Walker (2018) – rates of adaptive and non-adaptive evolution are lower between more dissimilar amino acids. It seems likely that these correlations are due to the mutational steps being smaller and hence that adaptive evolution proceeds via small steps in this component of evolution. Chen et al. (2019) apparently came to a different conclusion, but their analysis largely focussed on a statistic that is related to the proportion of substitutions that are adaptive, and hence conflates the pattern of adaptive and non-adaptive evolution. In fact, consistent with their findings and those of Bergman and Eyre-Walker (2018), we find the proportion of substitutions that are adaptive is uncorrelated to either the difference in volume or polarity (volume: r=-0.012, p=0.707; polarity: r=0.0003, p=0.314).

We demonstrate that population size change can lead to an artefactual correlation between a measure of adaptive evolution and any variable related to the mean strength of selection against deleterious mutations. For example, the negative correlation between *ω*_*a*_ and amino acid dissimilarity in *Drosophila* species could be generated by population expansion. It is therefore important to check whether the variable being considered is correlated to the mean strength of selection and/or confirm the correlation in a species which has undergone contraction.

## Materials and methods

### Data

We obtained gene sequences from Ensembl’s biomart (Yates et al. 2020) for the human GRCh38 genome build and for the Pan_tro_3.0 chimpanzee genome build. Orthologous genes were aligned using MUSCLE (Edgar, 2004). After filtering out genes with gaps that were not multiples of three we were left with 16,344 pairwise alignments. Numbers of synonymous and non-synonymous substitutions per site were obtained using PAML’s codeml (Yang, 2007) program. We used polymorphism data from the African superpopulation of the 1000 genomes dataset (The 1000 Genomes Consortium, 2015) to construct our site frequency spectra, with rates of adaptive and non-adaptive evolution estimated using Grapes (Galtier, 2016), under the “GammaZero” model. We chose African data because the African population is thought to have undergone less complex demographic changes then other human populations (Gutenkunst et al. 2009; Gravel et al. 2011). We fitted a weighted regression to our estimates of the rate of evolution, weighting by the reciprocal of the variance for each estimate of ω_a_ and ω_na_ . The confidence interval and variance on our estimates of ω_a_ and ω_na_ were obtained by bootstrapping the dataset by gene 100 times.

### RSA analysis

In order to obtain structural information for each protein sequence, we ran blastp (Schaffer, 2001) to assign each protein sequence to a PDB structure, and respective chain, by using the “pdbaa” library and an *E*-value threshold of 10^−10^. In instances of multiple matches, the match with the lowest *E*-value was kept. The corresponding PDB structures were further processed to only keep the corresponding chain per polymer. PDB manipulation and analysis were carried on using the R package “bio3d” (Grant et al. 2006). Values for solvent accessibility (SA) per residue were obtained using the “dssp” program with default options. To map SA values to each residue of the protein sequence a pairwise alignment between each protein and the respective PDB sequence was performed with MAFFT, allowing gaps in both sequences in order to increase the block size of sites aligned. The final data set comprised a total of 7,984,041 sites with SA information. We computed the RSA by dividing SA by the amino-acid’s solvent accessible area (Tien et al. 2013), giving us a final dataset of 3,505,615 sites for which we have RSA information.

These sites were grouped into 20 RSA bins of roughly equal size in terms of the number of sites, with rates of adaptive and non-adaptive evolution estimated for each bin. These rates were correlated with the mean RSA of each bin.

### Amino acid dissimilarity analysis

For the amino acid dissimilarity analysis we followed the methodology outlined in Bergman and Eyre-Walker (2019), with amino acid polarity and volume scores taken from data available in the AAindex1 database (Kawashima et al. 2008). We compared the SFS for a particular amino acid pair with synonymous data from 4-fold degenerate codons separated by the same mutational step. For example, alanine and glycine are separated by a single nucleotide change (C<>G at second position). Therefore, we used the SFS and divergence for all 4-fold degenerate codons separated by a single C<>G mutational step in estimating ω_a_ and ω_na_ . For amino acids separated by more than one mutational step (e.g. a C<>G or an A<>T mutational step), we used a weighted average SFS from the SFSs for the mutational types at 4-fold sites, weighting by the frequency of the respective codons as in Bergman and Eyre-Walker (2019).

For the analysis involving p_N_ /p_S_ we used a hypergeometric distribution to resample the SFS, and generate two SFSs, one used to estimate rates of adaptive and non-adaptive evolution, and one used to estimate p_N_ /p_S_ .

## Notes

### Competing Interest Statement

The authors have declared no competing interest.

