## supplementary material for "Site level factors that affect the rate of adaptive evolution in humans and chimpanzees; the effect of contracting population size"

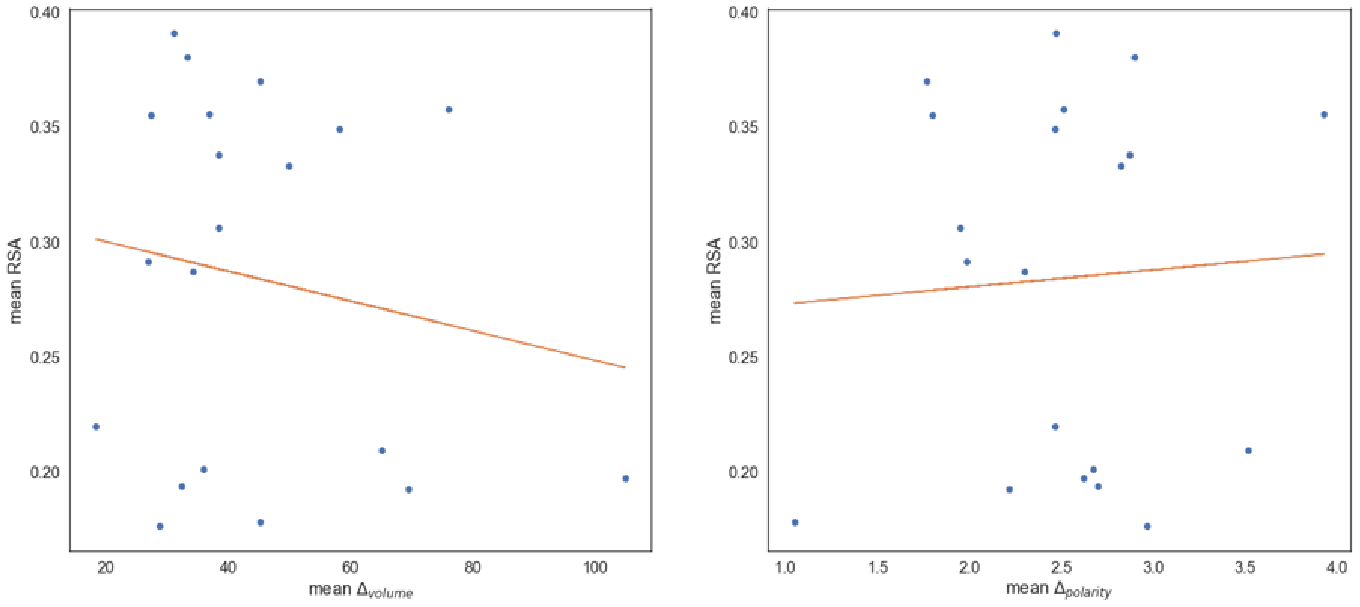


**Supplementary figure S1:** Average RSA of an amino acid and the average difference in volume or polarity to its one mutation step neighbours.

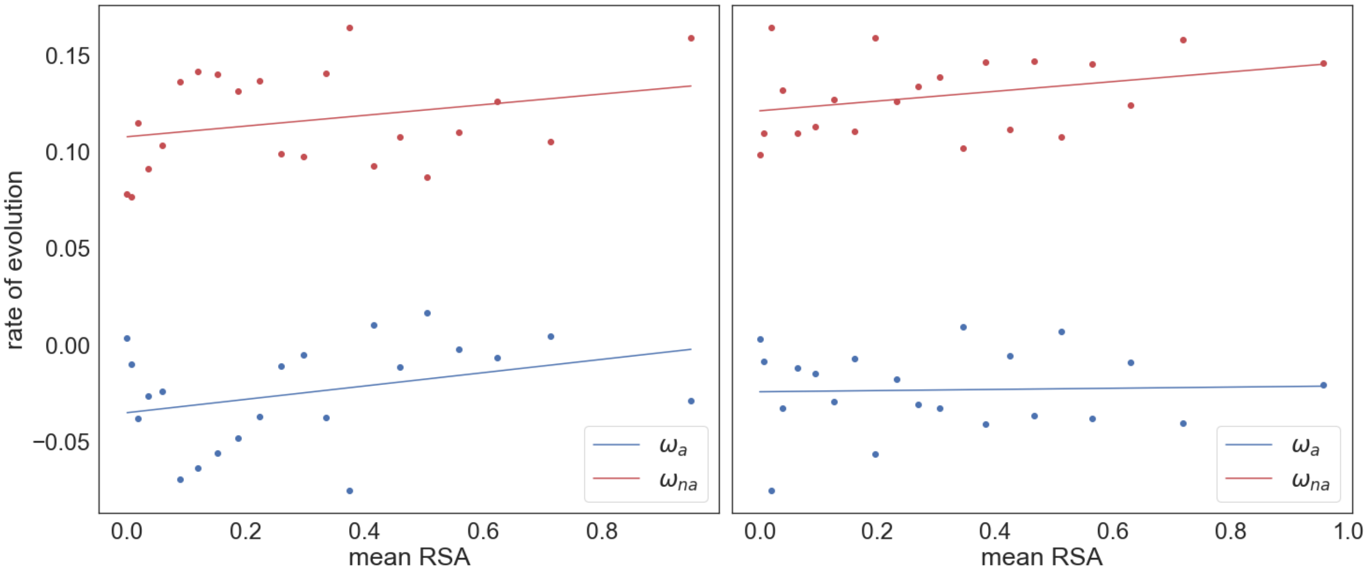


**Supplementary Figure S2:** Estimates of $\omega_a$ and $\omega_{na}$ plotted against mean relative solvent accessibility, controlling for volume difference (left) and polarity difference (right). Data binned into 20 RSA bins of roughly equal size. For each analysis, a weighted linear regression is fitted to the data. The respective significance of each correlation is shown in the plot legend, (*$P < 0.05$; **$P < 0.01$; ***$P < 0.001$; "." $0.05 \leq P < 0.10$) for $\omega_a$ and $\omega_{na}$). Regression is weighted by the reciprocal of the variance for each estimate of $\omega_a$ and $\omega_{na}$, which were estimated by bootstrapping the data by gene 100 times for each data point.

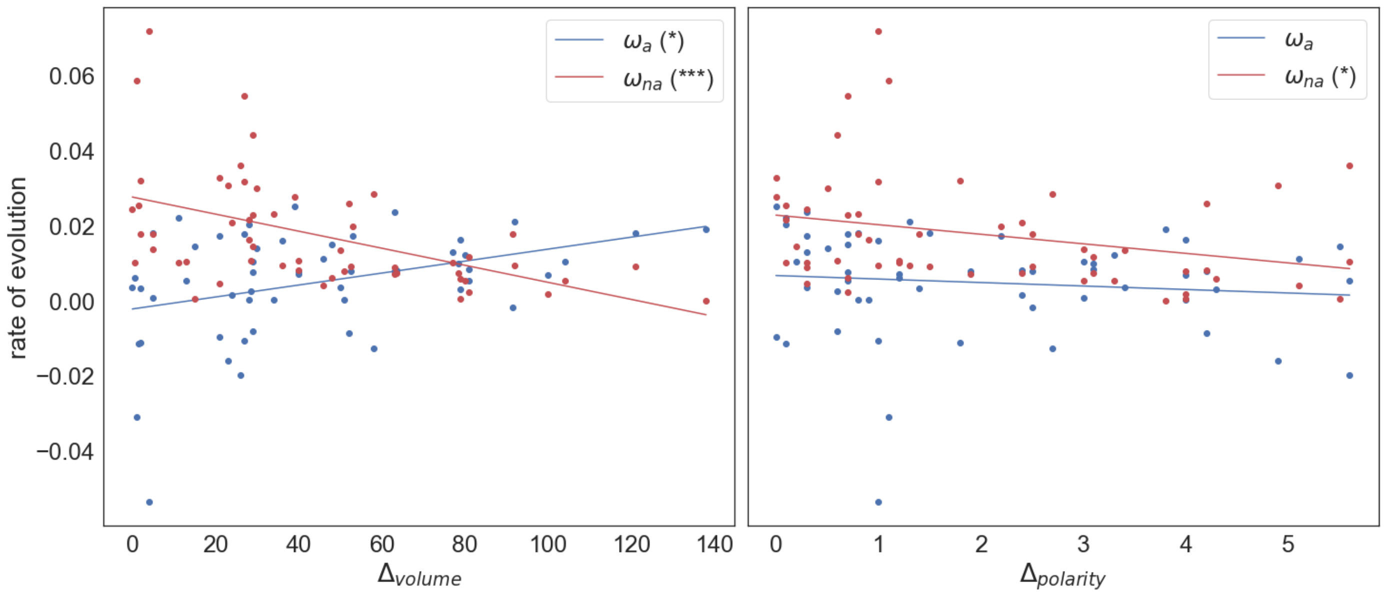


**Supplementary Figure S3:** The adaptive and non-adaptive substitution rate plotted against the difference in a) volume, b) polarity, controlling for relative solvent accessibility. A weighted linear regression is fitted to the data, weighted by the variance of each estimate. The respective significance of each correlation is shown in the legend, (*$P < 0.05$; **$P < 0.01$; ***$P < 0.001$; "." $0.05 \leq P < 0.10$).
